# VLIB: Unveiling insights through Visual and Linguistic Integration of Biorxiv data relevant to cancer via Multimodal Large Language Model

**DOI:** 10.1101/2023.10.31.565037

**Authors:** Vignesh Prabhakar, Kai Liu

## Abstract

The field of cancer research has greatly benefited from the wealth of new knowledge provided by research articles and preprints on platforms like Biorxiv. This study investigates the role of scientific figures and their accompanying captions in enhancing our comprehension of cancer. Leveraging the capabilities of Multimodal Large Language Models (MLLMs), we conduct a comprehensive analysis of both visual and linguistic data in biomedical literature. Our work introduces VLIB, a substantial scientific figure-caption dataset generated from cancer biology papers on Biorxiv. After thorough preprocessing, which includes figure-caption pair extraction, sub-figure identification, and text normalization, VLIB comprises over 500,000 figures from more than 70,000 papers, each accompanied by relevant captions. We fine-tune baseline MLLMs using our VLIB dataset for downstream vision-language tasks, such as image captioning and visual question answering (VQA), to assess their performance. Our experimental results underscore the vital role played by scientific figures, including molecular structures, histological images, and data visualizations, in conjunction with their captions, in facilitating knowledge translation through MLLMs. Specifically, we achieved a ROUGE score of 0.66 for VQA and 0.68 for image captioning, as well as a BLEU score of 0.72 for VQA and 0.70 for image captioning. Furthermore, our investigation highlights the potential of MLLMs to bridge the gap between artificial intelligence and domain experts in the field of cancer biology.

## Introduction

The field of cancer research constantly evolves with the dissemination of new knowledge through research articles and preprints, often accessible via platforms like Biorxiv [1]. This repository of preprints not only broadens our comprehension of cancer through written descriptions but also employs visual aids such as molecular structures, histological images, data visualizations, and other scientific figures. This study investigates the synergy between these scientific figures and their accompanying captions to harness their combined potential in a multimodal learning framework, enriching our understanding of contemporary cancer research [2].

By leveraging the capabilities of Multimodal Large Language Models (MLLMs), we delve into the fusion of visual and linguistic narratives within the biomedical literature [3]. This convergence facilitates a more comprehensive grasp of the domain knowledge inherent in the subject matter. To this end, we introduce VLIB, an extensive repository of scientific figure-caption pairs curated from a subset of cancer biology research papers featured on Biorxiv. The creation of VLIB involved rigorous preprocessing steps, including the extraction of figure-caption pairs, the identification of sub-figures within a broader visual context, and text normalization [4]. This meticulously assembled collection encompasses over half a million figures sourced from more than 70,000 papers, each paired with its corresponding natural language caption.

Our methodology involves assessing the performance of baseline MLLMs in the domain of vision-language tasks, encompassing image captioning and visual question answering (VQA) [5][6]. Our experiments delve into the intricate interplay between vision and language, shedding light on the opportunities and challenges associated with generating captions for scientific figures. Our analysis underscores the indispensable role played by a diverse array of scientific figures—ranging from molecular structures to histological images and data visualizations—in conjunction with their accompanying captions. This harmonious interplay bridges two modalities of data, enabling the translation of complex scientific concepts into accessible knowledge through the capabilities of MLLMs [3].

Furthermore, our exploration extends to the potential of MLLMs to bridge the gap between artificial intelligence and domain expertise in the expansive field of cancer biology. As our study unfolds, the interaction between MLLMs and subject matter experts emerges as a compelling narrative, hinting at the profound potential of collaborative synergies to unlock new dimensions of understanding within the realm of cancer research.

### Related Work

*SciCap* introduces an innovative framework for automatically generating high-quality captions for scientific figures within research articles [4]. It emphasizes the critical role of informative figure captions in effectively communicating complex data and addresses the common issue of inadequate captions. The authors curated the significant SCICAP dataset, comprising over two million figures extracted from more than 290,000 computer science arXiv papers published between 2010 and 2020 [7]. Through meticulous preprocessing, including figure-type classification, sub-figure identification, and text normalization, this dataset was prepared with dual objectives: to aid researchers in crafting improved figure captions and to enhance accessibility for visually impaired readers. This work lays the foundation for automatic figure-captioning models, catering specifically to scientific figures, an area that has received limited attention thus far. The study provides insight into both promising opportunities and complex challenges associated with generating captions for scientific figures, as demonstrated through baseline model experiments.

The *ScienceQA* dataset addresses the limitations of existing science question benchmarks by introducing a comprehensive and multimodal benchmark containing approximately 21,208 diverse science-related multiple-choice questions [8]. These questions span various science subjects, including natural science, language science, and social science, accompanied by annotations for answers supported by corresponding lectures and explanations. With a focus on enabling multi-hop reasoning, *ScienceQA* serves as a valuable resource for evaluating the multi-modal reasoning capabilities of AI models, shedding light on how deep learning models, including large-scale language models, synthesize chains of thought to arrive at answers [9]. The dataset’s inclusion of lectures and explanations associated with answers contributes to the interpretability and transparency of AI reasoning processes, highlighting its novelty and utility in advancing AI research within this domain.

The introduction of the *Multimodal C4 (MMC4)* dataset addresses the need for publicly available large-scale data that supports in-context learning for vision and language models. This multimodal dataset extends the text-only *c4* corpus, incorporating image-text sequences interleaved in a manner that facilitates both few-shot learning and complex prompts involving interactions between images [10]. An algorithm based on CLIP features effectively aligns images within longer text segments. The authors demonstrate the relevance and alignment of images with associated text, along with the successful application of filters to ensure appropriateness and accuracy. The dataset opens up possibilities for various applications, including an open-source version of the Flamingo model [11], advancing the availability of multimodal data for in-context learning and enhancing performance in tasks like image captioning through the use of the MMC4 dataset.

The *LAION-5B* dataset addresses the need for publicly available large-scale image-text data, particularly for training state-of-the-art language-vision models like CLIP and DALL-E [12]. While these models exhibit strong capabilities in text-guided image generation and zero-shot classification, limited access to large-scale datasets hindered broader research in this field. *LAION-5B* offers a dataset comprising 5.85 billion CLIP-filtered image-text pairs, enabling the replication and fine-tuning of foundational models and further exploration of multimodal model capabilities. This effort underscores the importance of democratizing research by providing such a dataset, filling a gap in publicly available resources for the broader research community. While positioned as a crucial step toward large-scale multimodal pre-training, the dataset’s technical limitations and ethical considerations are also discussed, emphasizing the evolving nature of research in this domain.

The *Med-Flamingo* paper addresses the challenges of medical domain applications, where data availability is often limited, by introducing a novel multimodal few-shot learner. Building upon the *OpenFlamingo-9B* model, the authors extend pre-training with paired and interleaved medical image-text data from PMC-Open Access publications and textbooks [13], resulting in *Med-Flamingo*. This model enables few-shot generative medical visual question answering (VQA) capabilities and is evaluated on various datasets, including a new open-ended VQA dataset of visual USMLE-style problems. Notably, the paper introduces a human evaluation, where physicians assess generative medical VQA in an interactive application. *Med-Flamingo* demonstrates improvements of up to 20% in clinician ratings for generative medical VQA, showcasing its potential in enabling multimodal medical few-shot adaptations and highlighting its contributions to enhancing medical applications.

*LLaVA-Med* addresses the limitations of applying multimodal conversational AI models to the biomedical domain, given their reliance on general-domain image-text pairs that inadequately capture the complexities of biomedical information [14]. To overcome this challenge, the authors introduce a novel approach centered around the Large Language and Vision Assistant for BioMedicine (*LLaVA-Med*). This approach involves leveraging a comprehensive biomedical figure-caption dataset from PubMed Central, employing GPT-4 to generate diverse instruction-following data [15]. Through fine-tuning this model using a curriculum learning strategy, *LLaVA-Med* showcases impressive proficiency in responding to open-ended biomedical image-related inquiries. The contribution extends to a new data generation pipeline, the *LLaVA-Med* model, and insights into domain-specific multimodal conversational assistant adaptation [16]. By sharing the biomedical instruction-following dataset and codebase, the authors foster progress in biomedical multimodal learning research. This work underscores the potential of adapting multimodal conversational AI to intricate verticals and establishes a significant advancement in the realm of biomedical dialogue systems.

The *PathVQA* dataset represents a pioneering effort to address the challenges of generating a visual question answering (VQA) dataset for medical pathology, particularly aligned with the requirements of the American Board of Pathology (ABP) certification examination [17]. Distinguished by its focus on the medical domain, the dataset’s creation involves a strategic combination of pathology textbooks and digital resources to overcome the inherent complexities and limitations of medical image understanding. Through a semi-automated pipeline, the dataset incorporates pathology images, precise captions, and question-answer pairs generated from these resources. Comprising 4,998 pathology images and 32,799 associated question-answer pairs, the dataset holds promise as a critical resource for advancing research in medical VQA. With a thorough overview of existing VQA datasets and the application of established and state-of-the-art VQA methods, the authors contribute a comprehensive framework to facilitate benchmarking and innovation in the realm of medical VQA [5]. This initiative not only establishes a valuable dataset but also lays the foundation for future advancements in AI-assisted clinical decision-making and medical education.

The *ARCH* dataset emerges as a significant contribution to the field of computational pathology (CP), offering a novel perspective on data representation and dense supervision for CP tasks. Distinct from existing CP datasets, *ARCH* presents an extensive array of diagnostic and morphological descriptions across various stains, tissue types, and pathologies [18]. Through a comprehensive dataset of dense image captions obtained from clinical and academic pathology sources, the authors postulate that pre-trained encoders from these captions could yield transferable and universal CP representations. This conjecture is supported by the demonstration that *ARCH* representations outperform ImageNet features or representations achieved through self-supervised or multi-task learning. By drawing parallels between CP sub-tasks and image captioning, and substantiated with attention plots, the paper suggests that the intricate relationships among CP sub-tasks embedded within captions result in more general CP image features. Overall, *ARCH* not only introduces a valuable dataset but also offers a potential avenue for addressing the challenges of information-rich ground truth annotation in medical imaging.

The OpenPath V2 dataset represents a groundbreaking effort to address the scarcity of annotated medical images for advancements in computational pathology [19]. This dataset harnesses the collective knowledge shared by clinicians on public platforms like medical Twitter, comprising 208,414 pathology images paired with natural language descriptions. It stands as the most extensive publicly available collection of pathology images annotated with textual descriptions. The significance of this resource becomes evident through the development of the PLIP (Pathology Language-Image Pre-training) model, trained on OpenPath. Through this model, state-of-the-art zero-shot and transfer learning capabilities are achieved for classifying diverse pathology images. Moreover, PLIP facilitates the retrieval of similar cases through both image and natural language searches, promising a significant contribution to knowledge sharing within the pathology community. This study underscores the potential of publicly shared medical information as a valuable resource to propel the field of biomedical AI forward.

In summary, the reviewed literature presents a spectrum of datasets and models, each addressing unique challenges and contributing significantly to advancing multimodal learning, particularly within specialized domains such as medicine and computational pathology. These resources collectively represent a rich tapestry of efforts, from improving figure-captioning in scientific articles to enabling multimodal reasoning in AI models. As we delve deeper into the intersection of vision and language, these datasets and models serve as pivotal assets, paving the way for innovations in healthcare, scientific research, and AI-assisted decision-making. Their availability not only fosters progress within specific domains but also sets the stage for interdisciplinary collaborations, emphasizing the transformative potential of multimodal learning in our evolving technological landscape.

## Constructing the VLIB Dataset

### Data Acquisition and Preprocessing

Our dataset represents a pioneering endeavor in the aggregation and organization of information sourced from a multitude of cancer-related papers available on BioRxiv. Utilizing the BioRxiv API, this dataset compiles a diverse collection of scientific articles that delve into various facets of cancer research [1]. What sets our dataset apart is its multimodal nature, which combines textual insights with visual data, drawing from a vast and open-access biomedical literature corpus.

### Figure-Caption Pair Extraction

To enhance the depth and breadth of information captured, our dataset utilizes Transformer-based Optical Character Recognition (OCR) to extract and preserve images embedded within these papers [20]. Additionally, we employ web scraping techniques with the aid of an HTML parser to meticulously gather relevant text content from the HTML representations of the articles [21]. This integrated approach ensures the accessibility of both structured and unstructured information, laying the foundation for a comprehensive multimodal vision-language dataset.

### Text Normalization

To ensure consistency and facilitate analysis, we employed various standard text normalization techniques for the textual content associated with the images extracted from BioRxiv. These techniques encompass stemming, lemmatization, the incorporation of custom vocabularies and ontologies to handle domain-specific terminologies, and the enforcement of standardized naming conventions for genes and proteins where required [22].

### Data Summary

The VLIB dataset represents an extensive compilation, encompassing roughly 500,000 figures sourced from a vast corpus of over 70,000 research papers. Each figure within the dataset is meticulously paired with its corresponding caption, offering essential context and insights into the visual content.

## Experimental Results

### Baseline Model

In this study, we harnessed the capabilities of BEiT-3, a cutting-edge vision-language model built upon the foundations of Multiway Transformers [23][24]. This architecture is meticulously designed to excel in encoding and seamlessly integrating different modalities, making it particularly well-suited for a plethora of multimodal understanding tasks, encompassing both vision and language. The core of BEiT v3 is constituted by Multiway Transformer blocks, each featuring a shared self-attention module and modality-specific experts, as illustrated in Figure 3. This integration facilitates efficient alignment between modalities and deep fusion of information, yielding outstanding performance across tasks such as image classification, object detection, instance segmentation, and semantic segmentation.

One of the model’s paramount strengths lies in its versatility, serving as a robust image backbone for a multitude of vision-related tasks, thus significantly enhancing its utility in the realm of computer vision. Furthermore, BEiT v3 is amenable to fine-tuning for tasks such as image-text retrieval, where it operates as a dual encoder, or for a diverse range of multimodal understanding and generation tasks, courtesy of its exceptional multimodal capabilities.

The pretraining approach for BEiT-3 is underpinned by a unified mask-then-predict methodology, imparting the model with the capacity to learn high-quality representations while capturing intricate alignments between different modalities, as depicted in Figure 2. This involves tokenization of text data using SentencePiece and image data using the BEiT V2 tokenizer, allowing for the extraction of discrete visual tokens [23][25]. BEiT-3’s pretraining data comprises an extensive dataset, encompassing millions of images and image-text pairs sourced from diverse public datasets, alongside substantial monomodal data from ImageNet-21K and an array of diverse text corpora [26]. In essence, BEiT v3 represents a monumental leap in vision-language models, delivering a versatile and potent solution for a broad spectrum of tasks, spanning from intricate image processing to intricate multimodal understanding and generation, thus setting new benchmarks within the field of multimodal AI.

**Figure 1.**
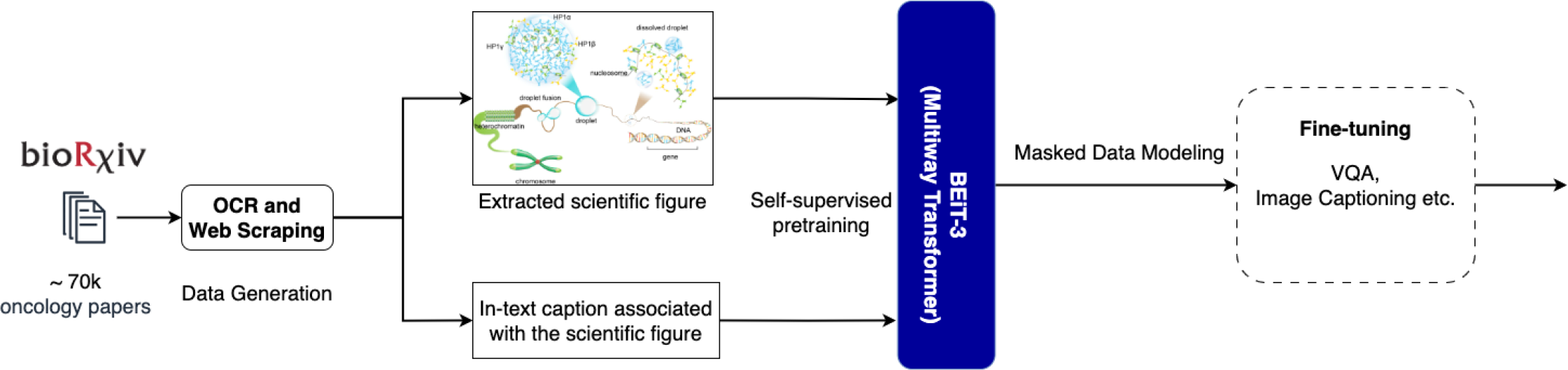
End to end architecture of VLIB dataset construction and vision-language modeling using BEiT-3

**Figure 2.**
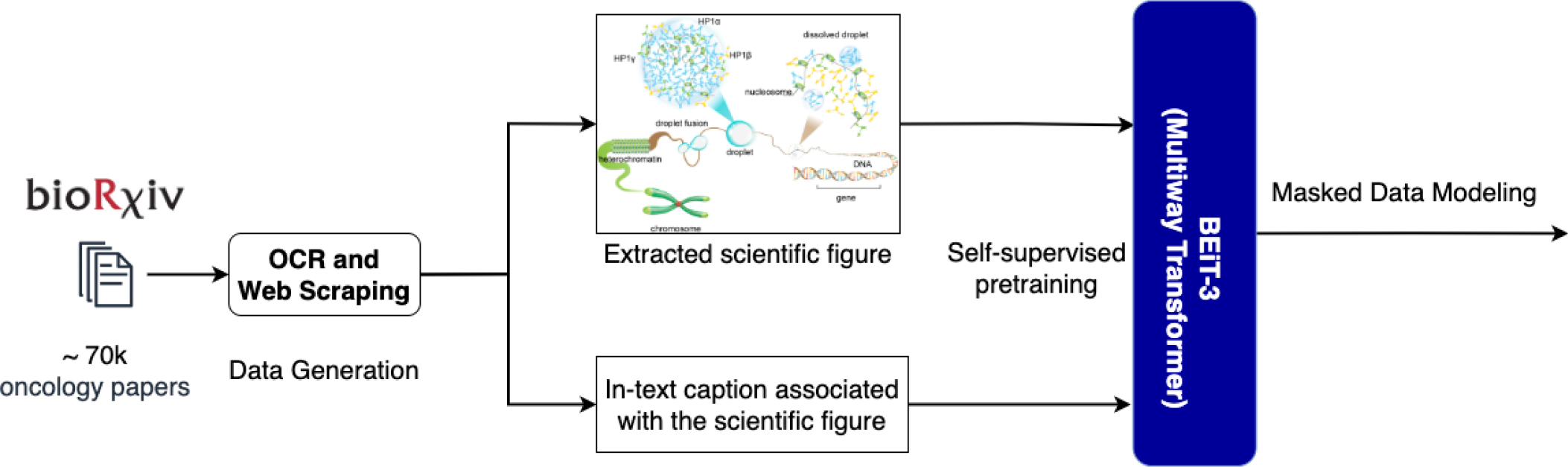
Self supervised pretraining of BEiT-3 using figures and relevant text extracted from Biorxiv oncology papers

**Figure 3.**
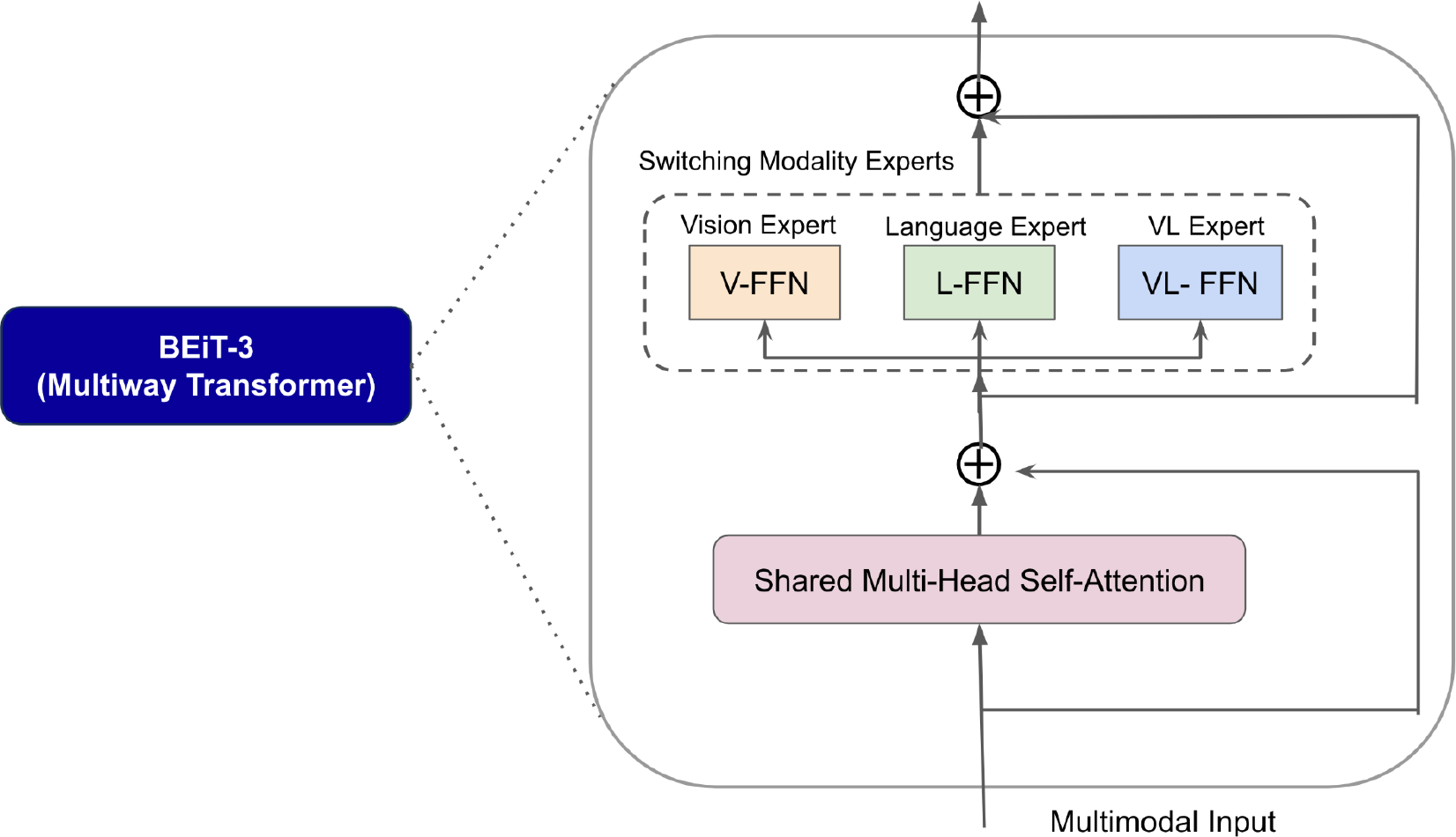
Architecture of the multiway transformer of BEiT-3 that accepts multimodal input to project multimodal vector representations to a shared latent space.

### Experiment Setup

In this section, we detail the experimental setup employed to leverage the BEiT-3 vision-language model. Our goal was to perform self-supervised pretraining on the BioRxiv image-text pairs, followed by fine-tuning the model for visual question answering (VQA) and image captioning tasks [23][27]. The overarching objective was to enhance the model’s understanding of the intricate relationships between images and text within the biomedical domain.

We initiated the process by utilizing the BEiT-3 vision-language model for self-supervised pretraining, as depicted in Figure 2. The model underwent masked data modeling during pretraining, wherein a portion of input tokens (both image and text modalities) were randomly masked, prompting the model to predict these masked tokens [28]. This pretraining phase equipped the model with a robust comprehension of the joint distribution between images and their corresponding textual descriptions.

Subsequently, the fine-tuning phase began. The BEiT-3 model was fine-tuned on two specific downstream tasks: visual question answering (VQA) and image captioning [5][6]. For VQA, we generated question-answer pairs using another multimodal large language model (MLLM) called MiniGPT-4 [29]. These questions pertained to details within the images, with answers generated by MiniGPT-4 serving as ground-truth responses. In the image captioning task, the model learned to generate relevant captions for the input images, as illustrated in Figure 5.

**Figure 4.**
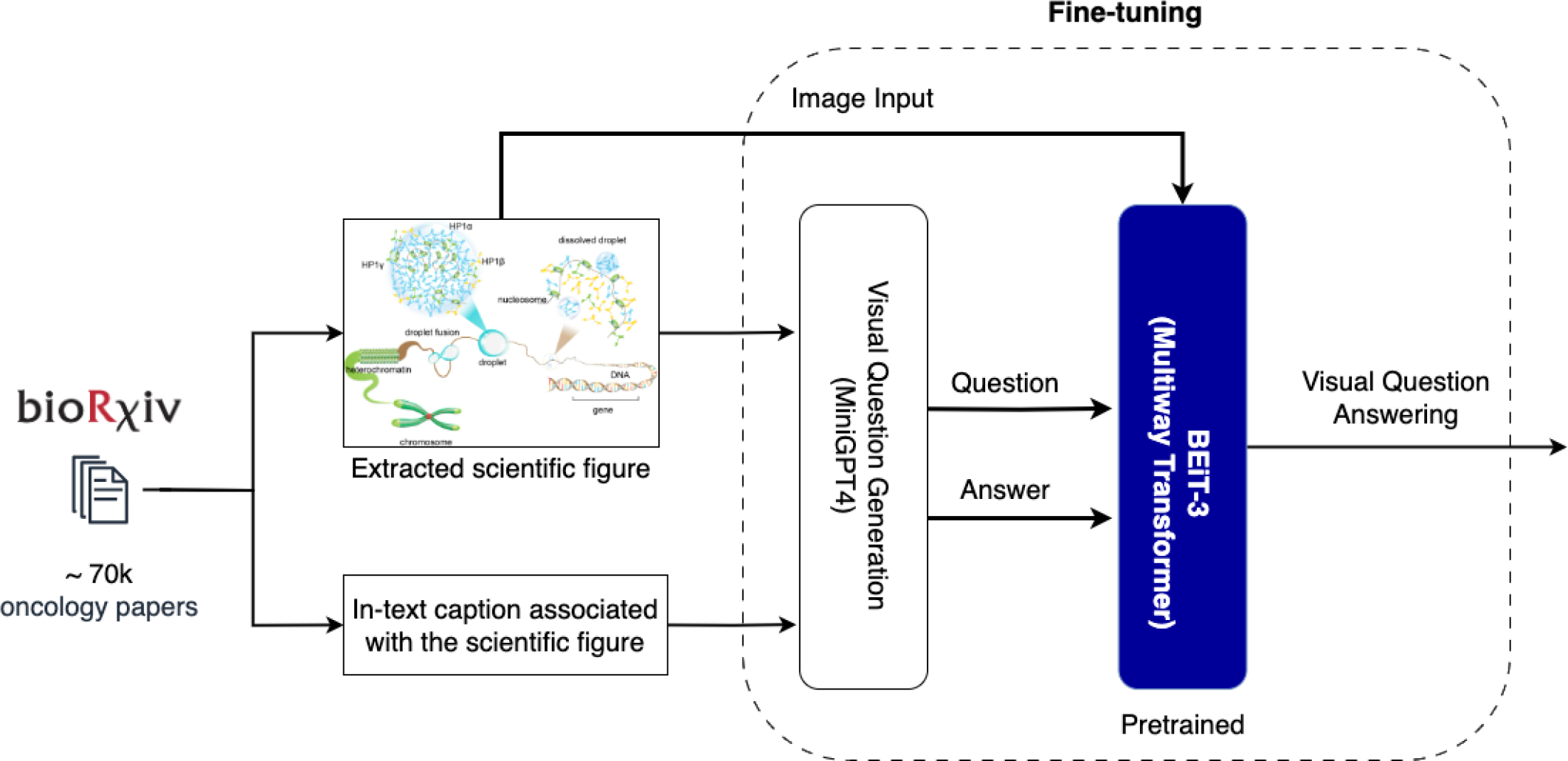
Fine Tuning BEiT-3 on Visual Question Answering task (VQA) using question, answer pairs relevant to the Biorxiv figures and text generated by MiniGPT-4

**Figure 5.**
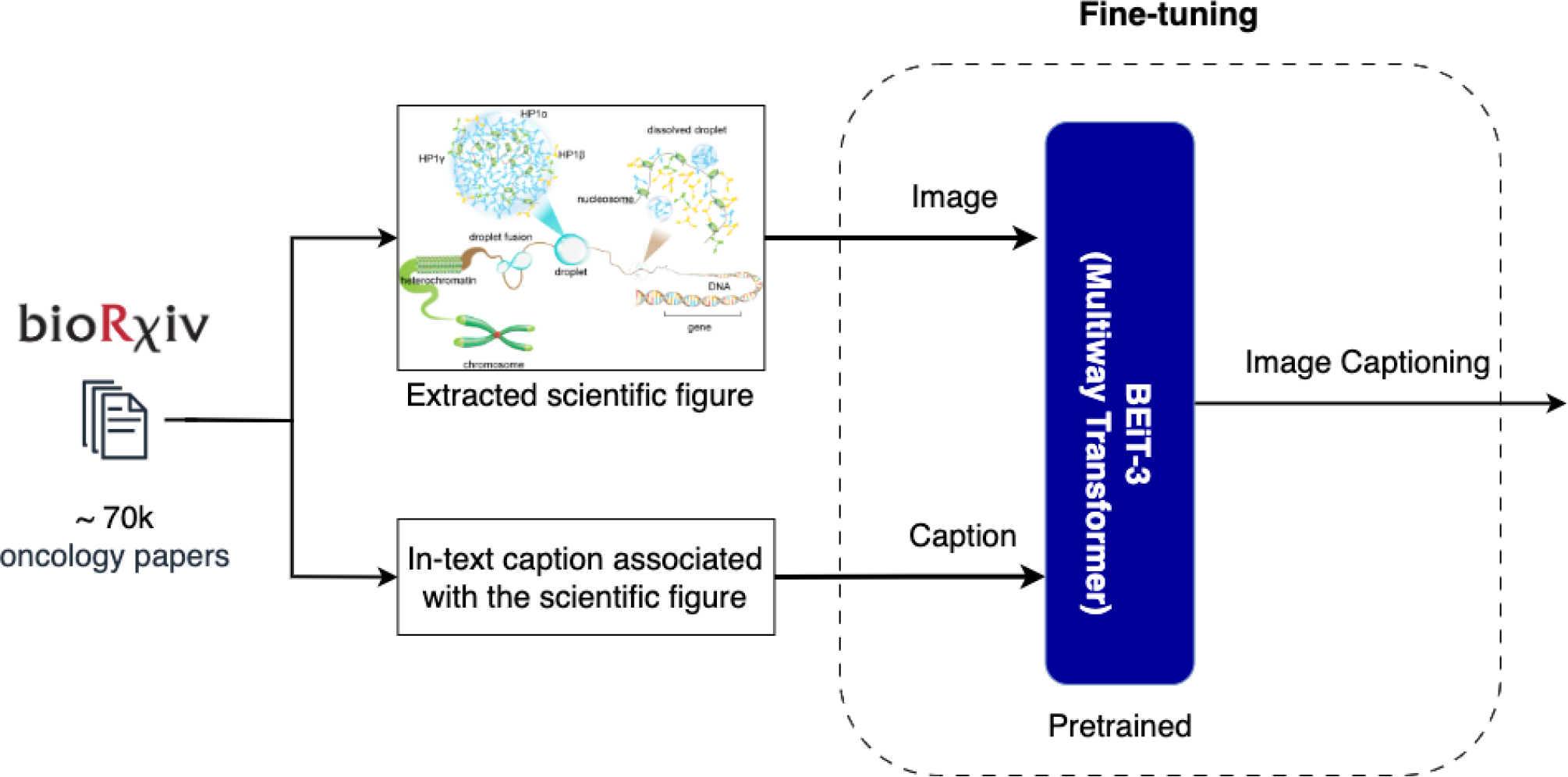
Fine Tuning BEiT-3 on Image Captioning task using scientific figures and their respective captions extracted from Biorxiv

To support these experiments, we utilized 8 Nvidia V-100 GPUs. Pretraining was conducted over 200 epochs to ensure comprehensive convergence and robust feature extraction. Leveraging 8 GPUs allowed for efficient parallel processing, significantly expediting the training process.

Our experiment setup aimed to lay a solid foundation for the BEiT-3 model’s multimodal understanding of biomedical images and text. The model’s performance in downstream tasks, such as VQA and image captioning, serves as a testament to its capacity to bridge the gap between visual and textual information within the intricate context of biomedical research.

### Evaluation and Results

To assess the quality of machine-generated text, especially in tasks such as machine translation, we employ the BLEU metric. BLEU measures the similarity between a reference (human-generated) translation and an automatic (machine-generated) translation [30]. It quantifies precision-based n-gram overlap between the candidate and reference translations. A higher BLEU score signifies a better match between candidate and reference translations. Additionally, we utilize the ROUGE metric family for evaluating tasks like automatic summarization, machine translation, and other natural language processing endeavors [31]. ROUGE calculates multiple measures, including precision, recall, and F1 score, based on the overlap of n-grams, word sequences, or even word-hyphen pairs between the machine-generated output and human reference summaries or translations.

In the context of Visual Question Answering (VQA) and Image Captioning, evaluating the natural language responses generated by the model is paramount to ensure the provision of relevant and coherent answers to questions about visual content [32]. Both BLEU and ROUGE scores are valuable tools for this purpose.

Although BLEU is primarily tailored for machine translation, it can be adapted effectively for VQA and Image Captioning tasks [32]. In VQA, the reference text is the ground truth answer provided by MiniGPT4, while the candidate text is the answer generated by our model fine-tuned for VQA. Similarly, in Image Captioning, the ground truth text is the figure caption extracted from the research paper, and the candidate text is the caption generated by our model fine-tuned for the image captioning task. BLEU assesses the alignment of n-grams and sequences in the model-generated answer with those in the human-provided answer, quantifying the degree to which the model’s response aligns with the expected answer. A higher BLEU score signifies a closer match and higher quality in the generated response.

ROUGE scores can also be applied to assess the quality of natural language responses from VQA and Image Captioning models [32]. Here, the reference answer/caption serves as the ground truth text, and the model’s answer is treated as the machine-generated text. ROUGE computes precision, recall, and F1 score based on n-gram overlap, aiding in quantifying how well the model’s response captures the essence of the actual ground truth. A high ROUGE score indicates that the model’s response is contextually and semantically similar to the expected answer.

In the VQA task, where the model must provide textual answers based on visual input and question prompts, BLEU and ROUGE scores provide valuable insights into the model’s language generation capabilities. These metrics assist in assessing the fluency, relevance, and correctness of the generated responses, ensuring that the VQA model delivers meaningful and coherent answers aligned with human expectations. Consequently, these scores serve as essential tools for fine-tuning and benchmarking VQA systems, ultimately enhancing their real-world performance.

As presented in Table 1, our model, pretrained on domain-specific scientific figures and text relevant to cancer biology from BioRxiv, achieved notable scores on the VQA and Image Captioning tasks. Specifically, on the VQA test set, our model achieved a BLEU score of 0.72 and a ROUGE score of 0.66. For the Image Captioning task, our model achieved a BLEU score of 0.70 and a ROUGE score of 0.68. These results highlight the model’s proficiency in generating contextually accurate responses and captions for visual content within the biomedical domain.

**Table 1:** The BLEU and ROUGE scores for VQA and image captioning downstream tasks performed on our VLIB dataset by BEiT-3.

|  | BLEU | ROUGE |
| --- | --- | --- |
| Visual Question Answering | 0.72 | 0.66 |
| Image Captioning | 0.70 | 0.68 |

## Conclusion and Future work

In conclusion, this paper introduced VLIB, a substantial image captioning dataset comprising scientific images and cancer-related captions sourced from BioRxiv. It sheds light on the dynamic interplay between scientific figures and their corresponding captions within the domain of cancer research, harnessing the formidable capabilities of Multimodal Large Language Models (MLLMs) to usher in a transformative era of understanding.

The birth of VLIB, a comprehensive repository housing over two million figure-caption pairs distilled from more than 70,000 cancer biology research papers on BioRxiv, serves as compelling evidence of the profound insights that emerge from the fusion of visual and linguistic narratives. Through meticulous preprocessing and rigorous experimentation, we’ve unveiled the pivotal role played by diverse scientific figures, ranging from intricate molecular structures to informative histological imagery. These figures act as catalysts, facilitating the translation of intricate domain knowledge into accessible insights, powered by the capabilities of MLLMs.

Importantly, our findings extend beyond the boundaries of artificial intelligence. They underscore the potential for MLLMs to foster collaborative exchanges between artificial intelligence and domain experts, thereby bridging critical gaps in the field of cancer biology research. By harnessing the synergies between these modalities, we envision a future where MLLMs emerge as invaluable tools for unlocking novel dimensions of comprehension and collaboration within the ever-evolving landscape of cancer research.

Looking ahead, our work opens up exciting avenues for future exploration. One promising direction involves enhancing the fine-tuning of MLLMs on domain-specific tasks within cancer research. Additionally, the continuous expansion and curation of VLIB can contribute to its role as a cornerstone dataset for advancing multimodal understanding in the biomedical domain.

Furthermore, the collaborative potential of MLLMs suggests the need for interdisciplinary research, fostering deeper connections between AI experts and domain specialists. This collaboration can lead to the development of specialized tools that aid in the interpretation of complex scientific data, ultimately accelerating breakthroughs in cancer biology.

As technology evolves and AI capabilities continue to advance, the synergy between visual and linguistic modalities promises to revolutionize how we perceive, interpret, and contribute to the vast body of knowledge in cancer research. Our work serves as a stepping stone towards this promising future, where the convergence of AI and domain expertise becomes a driving force for innovation and discovery.

## Ethical Considerations

### Data Licensing

The biorxiv dataset uses the Creative Commons Attribution (CC BY) license for the content shared on its platform. This license allows users to share, adapt and build upon the content from biorxiv research papers for any purpose (including commercial) as long as proper attribution is provided to the original authors and source. Furthermore by posting on biorxiv authors explicitly agree to text mining of their work.

### Absence of peer reviewed certification for the preprints

The biorxiv dataset consists of preprints that are not certified by scientific peer review. Therefore they could be prone to errors, conjectures, faulty reasoning and assumptions. Due to the very same reason the clinical accuracy of the responses derived from a MLLM trained on this dataset should be interpreted with a judicious sense of caution and scientific awareness.

## Acknowledgments

We would like to thank all the authors who published the preprint of their respective research articles on biorxiv. This work was supported and funded by Genentech Inc.

